# Microsatellite instability as a tool to diagnose genome instability in CHO cell culture

**DOI:** 10.1101/823252

**Authors:** Dylan G. Chitwood, Qinghua Wang, Kathryn Elliott, Aiyana Bullock, Dwon Jordana, Zhijang Li, Cathy Wu, Sarah W. Harcum, Christopher A. Saski

## Abstract

As bioprocess intensification has increased over the last 30 years, yields from mammalian cell processes have increased from 10’s of milligrams to over 10’s of grams per liter. Most of these gains in productivity have been due to increasing cell numbers in the bioreactors, and with those increases in cell numbers, strategies have been developed to minimize metabolite waste accumulation, such as lactate and ammonia. Unfortunately, cell growth cannot occur without some waste metabolite accumulation, as central metabolism is required to produce the biopharmaceutical. Inevitably, metabolic waste accumulation leads to decline and termination of the culture. While it is understood that the accumulation of these unwanted compounds imparts a less than optimal culture environment, little is known about the genotoxic properties and the influence of these compounds on global genome instability. In this study, we examined the effects on Chinese hamster ovary (CHO) cells’ genome sequences and physiology due to exposure to elevated ammonia levels. We identified genome-wide *de novo* mutations, in addition to variants in functional regions of certain genes involved in the mismatch repair (MMR) pathway, such as *DNA2, BRCA1* and *RAD52*, which led to loss-of-function and eventual genome instability. Additionally, we characterized the presence of microsatellites against the most recent Chinese Hamster genome assembly and discovered certain loci are not replicated faithfully in the presence of elevated ammonia, which represents microsatellite instability (MSI). Furthermore, we found 124 candidate loci that may be suitable biomarkers to gauge genome stability in CHO cultures.

## Introduction

Biopharmaceutical manufacturing represents nearly 2% of the total US GDP [1],thus an important driver of the US economy. Biopharmaceuticals include monoclonal antibodies, recombinant proteins, and assemblies of proteins produced by biological means. Commercial products are used as blood factors, thrombolytic agents, therapeutics, growth factors, interferons, and vaccines [2, 3].The most common mammalian cell line used is the CHO cell line, due to its ability to produce biopharmaceutical molecules with post-translational modifications required in humans [4]. However, it is well understood that recombinant CHO cell lines are susceptible to genome instability that is often observable after approximately 70 generations [5-8]. This genome instability reduces longevity in continuous culture; and productivity can be limited in fed-batch systems [9, 10]. A common occurrence in both continuous cultures and fed-batch systems is the accumulation of metabolic waste products, such as ammonia and lactate [11, 12]. The effects of these waste products on genome instability per se have not been directly assessed.

Microsatellite instability (MSI) is the observation of genetic hypermutability at microsatellite loci, where a high frequency of insertion or deletion (indel) mutations accumulate during cell division [13, 14]. Furthermore, the observation of MSI is evidence that the MMR machinery is not functioning properly [15]. Rather than correcting errors that occur spontaneously during DNA replication, cells with impaired MMR systems accumulate these errors globally, which are then passed on to daughter cells during division to become amplified as the culture persists. MSI loci are stable genetic biomarkers and have been utilized in the diagnosis of many cancers [16, 17]. Previous studies have shown that approximately 15% of patients with colorectal cancer [15, 18], 20% of patients with stomach cancer [19], and 30% of patients with endometrial cancer [20] could attribute their disease to genome instability that is diagnosed with microsatellite (MS) biomarkers. The clinical uptake of MSI-based diagnostics, such as the Bethesda Panel, demonstrates the reliability and clinical utility of MSI loci as biomarkers [21].

In this study, the effect of exposure to elevated levels of ammonia on genome instability during fed-batch cultures of CHO cells was investigated. Specifically, the accumulation of DNA mutations in cells exposed to elevated ammonia were compared to cultures grown under standard fed-batch conditions. Ammonia was added to duplicate parallel cultures at 10 mM and 30 mM final concentrations after 12 hours of culture time to establish a mild and high ammonia stress. After 72 hours of exposure to the elevated ammonia, samples were taken for genome sequencing and analysis of resulting SNP and indel variation. The SNPs and indels were mapped to the Chinese hamster genome, and assessed for functional impact in coding genes and presence in predicted microsatellite regions. The relevance of these findings advance our understanding of the genomic factors that lead to culture decline and establish a foundation for the development of DNA-based tools to imporove CHO cell biomanufacturing.

## Materials and Methods

### Culture Conditions

A recombinant CHO-K1 Clone A11 from the Vaccine Research Center at the National Institutes of Health (NIH), which expresses the anti-HIV antibody VRC01 (IgG_1_) was used. The inoculum train was expanded in 250 mL shake flasks with 70 mL ActiPro media (GE Healthcare) maintained at 5% CO_2_ and 37°C. The bioreactors were ambr®250 bioreactors (Sartorius Stedim, Göttingen, Germany) with two pitched blade impellers and an open pipe sparger (vessel part number: 001-5G25). The bioreactors were inoculated at a target cell density of 0.4 × 10^6^ cells/mL in ActiPro batch media and fed daily beginning on Day 3 (3% (v/v) Boost 7a and 0.3% (v/v) Boost 7b (GE Healthcare)). 12-h post inoculation, duplicate cultures were stressed with 0 mM, 10 mM, or 30 mM NH_4_Cl. The 0 mM cultures received equivalent volumes of water instead of the NH_4_Cl solution. Dissolved oxygen was controlled at 50% of air saturation using PID control that increased the O_2_ mixture in the gas sparge to 100%, then the stir speed from 300 to 600 rpm. 10% Antifoam Solution (SH30897.41 – GE Heathcare) was added as needed for foam control. For the ambr^®^ 250, all gases were supplied through the open pipe sparger. Overlay gassing was not used. The pH was controlled via sparging CO_2_ and air, and base pump (NaOH). The pH setpoint was 7.0 with a 0.2 deadband. Temperature was controlled at 37°C. Samples for MSI analysis were harvested at 84 hours culture time (72 hours post-stress) and centrifuged at approximately 2,000 x g for 15 minutes at 4°C. The supernatant was removed, and the pellet was stored at −80°C.

### DNA extraction, whole genome sequencing, and microsatellite variant discovery

Approximately 250 μg cell pellets were pre-washed with 1X phosphate buffered saline (PBS) prior to extraction. Total genomic DNA (gDNA) was purified with the DNAeasy Blood and Tissue Kit (Qiagen), following the manufacturer’s recommended procedures. Whole genome shotgun sequencing was performed on an Illumina NovaSeq (2×150 paired end) through a third-party vendor. Raw sequence data was assessed for quality with the FASTQC software (https://www.bioinformatics.babraham.ac.uk/projects/fastqc/). Raw sequence data was preprocessed to remove low quality bases and adapter sequences with the Trimmomatic software v.0.38 [22]. Preprocessed reads were aligned to the CriGri-PICR version assembly of the Chinese hamster genome (*Cricetulus griseus*) (RefSeq assembly accession: GCF_003668045.1) with the Bowtie2 v.2.3.4.1 short read aligner [23]. Alignments were coordinate sorted and indexed with SamTools v1.3.1 [24]. SNPs and indels were determined with the HaploTypeCaller Walker from the Genome Analysis Toolkit (GATK v.4.0) [25]. Functional SNPs were characterized with the SNPeffect software, v4.3 [26]. Genome-wide microsatellite loci were determined against the PICR CH assembly with MISA, a microsatellite finder software [27]. Microsatellite loci were intersected with indel coordinates using BedTools Intersect command 2.27.1 [28] to identify indel variants associated with microsatellites.

### Text/Data mining and Functional enrichment analysis

The query “genomic instability [MeSH Terms]” was used to search PubMed to retrieve the abstracts with PMIDs (13889 matched). The PubTator [29] tool was used to collect genes annotated in these abstracts with Entrez Gene IDs. Among the 5098 retrieved genes, 3054 genes were human (*Homo sapiens*), 845 genes were mouse (*Mus musculus*), representing the two largest groups of species. The ortholog pairs among human, mouse and Chinese hamster were mapped with NCBI ortholog assignment (ftp://ftp.ncbi.nlm.nih.gov/gene/DATA/gene_orthologs.gz). 15536 pairs of 1-to-1 human/Chinese hamster orthologs were obtained, which served as background for DAVID [30, 31] (http://david.abcc.ncifcrf.gov/) enrichment analysis. For the SNP lists with high, moderate, or low mutation effects, the Chinese hamster genes were mapped with corresponding human orthologs, then DAVID was used to obtain the enriched gene clusters, functional annotation clusters and charts for the given gene lists.

In other studies, environmental stressors, such as UV radiation [32], mutagenic compounds, and reactive oxygen species (ROS) [33], have demonstrated negative effects on genome integrity at the nucleotide level. For instance, ROS in tobacco smoke oxidize guanine bases to form 8-oxoguanine (8-oxoG), which results in the accumulation of single nucleotide polymorphisms (SNPs) [34]. SNP accumulation over time can lead to global genome instability in smokers, which is widely considered to be the leading cause of lung cancer in smokers [35]. Likewise, mutations, such as synonymous base changes in coding and regulatory regions, have little to no effect on gene transcription and translation. Yet, non-synonymous changes can have drastic effects on gene expression and cellular function. Specifically, alterations in critical genes that encode DNA MMR machinery can lead to global genome instability by failing to repair errors caused by a variety of events such as replication stalling [36], replication fork collapse [37, 38], double-strand breaks [39, 40], or environmental stressors.

### Identification of candidate MSI loci

The prevalence of indel mutations that intersect microsatellite loci are high in the ammonia stressed and control samples. In order to prioritize a list MSI loci with high confidence, a filtering strategy that leverages several criteria was established. First, each variable genomic locus was assigned a ‘mutation-score’ which is a proportion of the number of variant reads (allelic depth) by the total depth of reads for each site. Next, genomic sites were further ranked by ordering sites with the largest ‘mutation-score’ and then subtracting control samples from the treatment samples to identify sites where treated samples contained mutations and the controls did not, with sufficient depth of reads. The final ranked set of candidate MSI loci contain sites where control samples do not contain any variant reads, with sufficient read depth in each sample. The filtering method is summarized below in Fig 1.

**Fig 1:**
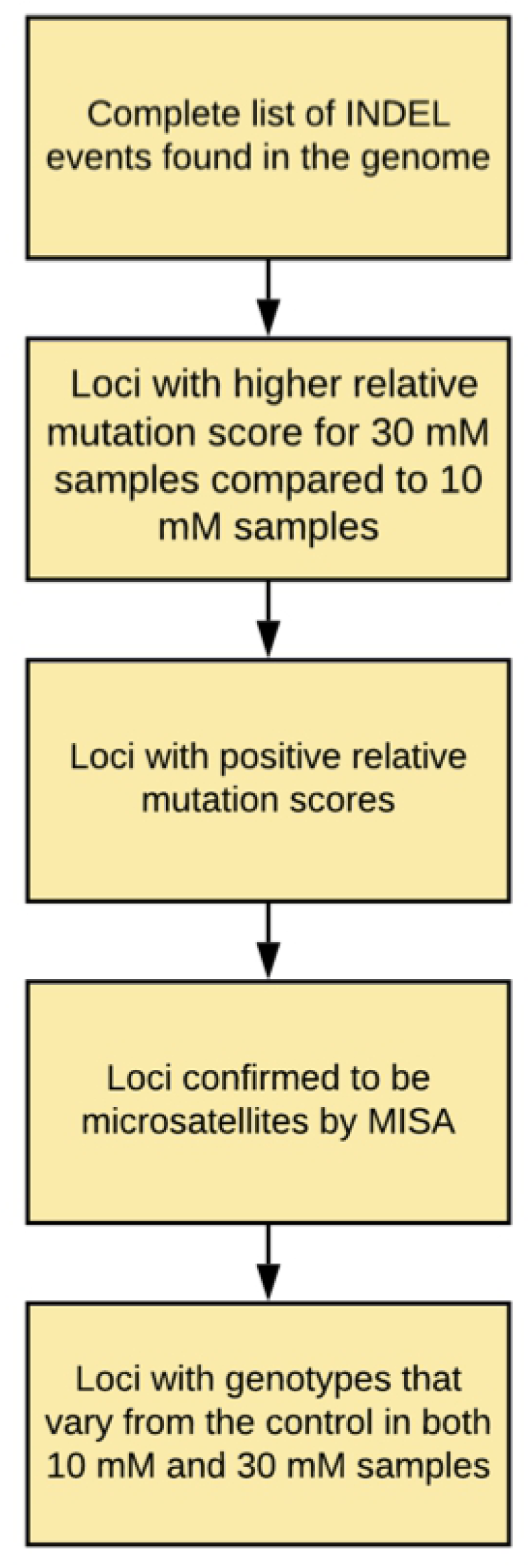
Progression of candidate loci through the multiple filtering criteria.

## Results

### Growth and metabolite profiles

CHO cells were cultured in highly controlled ambr^®^ 250 bioreactors for 12 hours prior to the addition of ammonia to stress the cultures. Up to 1.5 days, there was no observable difference in the viable cell density (VCD); however, by 2.5 days, the 30 mM ammonia-stressed cultures had substanially decreased VCD compared to the control and 10 mM stressed cultures (Fig 2A). The ammonia profiles are shown in Fig 2B. The 10 mM ammonia-stressed cultures had similar VCDs to the control cultures until Day 7; yet always had cell viabilities that were similar to the control cultures. In contrast, the 30 mM ammonia-stressed cultures reach peak VCD on Day 4 and the cultures were harvested on Day 8.5 due to low viability (< 70%). A cell viability below 70% is a standard harvesting threshold. The samples for genome sequencing were taken at 84 hours of culture time (Day 3.5), i.e. 72 hours post-stress. At the time of harvest of genome sequencing samples, the viability for all samples was greater than 90% (Fig 2A). The glucose and lactate profiles (Fig 2D,E) indicate that the control and 10 mM ammonia-stressed cultures had very similar metabolic characteristics throughout the entire cultures; however, the product titer was lower for the 10 mM ammonia-stressed cultures (Fig 2C). In contrast, the 30 mM ammonia-stressed cultures accumulated glucose and lactate after Day 5, due to the feeding protocol and the lack of cell growth. All cultures received the feed Boost 7a, which contains glucose, based on the culture volume and not the VCD. It is well-known that excessive glucose causes lactate accumulation [12]. Amino acid profiles were also obtained for these cultures. The most interesting amino acid profile was for alanine, where the 10 mM and 30 mM ammonia-stressed culture profiles were very similar through Day 6, while the control culture has a significantly different profile (Fig 2F). Otherwise, the control and 10 mM ammonia-stressed cultures had similar profiles, even if the concentrations were not identical. In summary, the samples for the genome sequencing analysis were taken when there were no substantial VCD, viability, or metabolic difference between the control and 10 mM ammonia-stressed cultures, and only the VCD for the 30 mM ammonia-stressed cultures was indicative of the stress (Fig 2).

**Fig 2:**
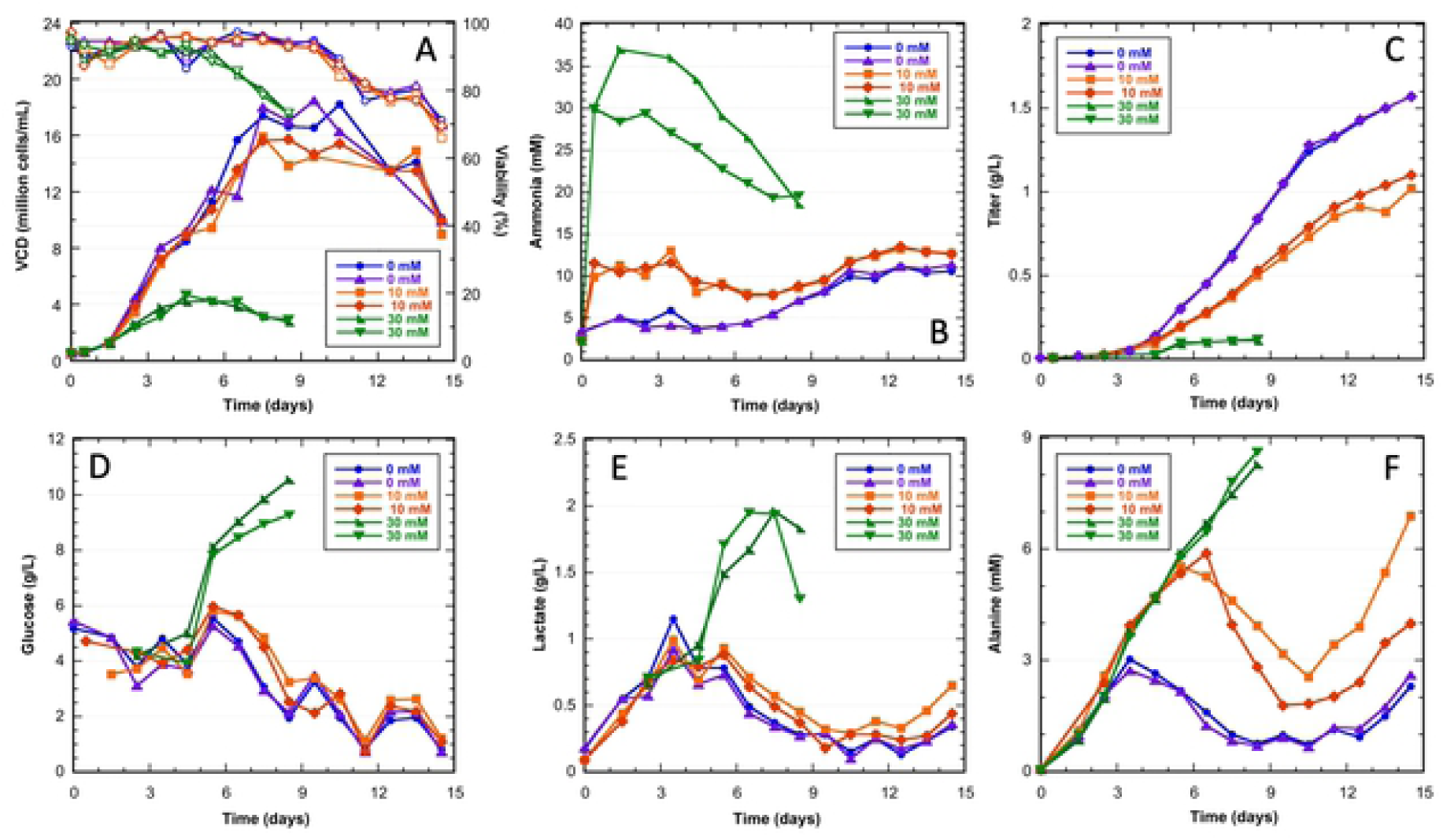
Growth, ammonia, titer and metabolic profiles for CHO VRC01 cell line cultured in duplicate in the ambr^®^250. The 10 mM and 30 mM ammonia stresses were added at 12 hours. A) Viable cell density (VCD) and viability (filled and hollow symbols, respectively), B) ammonia, C) titer, D) glucose, E) lactate, and F) alanine. Control - 0 mM (blue and purple lines); 10 mM ammonia (orange and red lines); 30 mM ammonia (green and dark green lines).

### Whole genome shotgun sequencing and variant discovery in stressed conditions

Whole genome shotgun sequences were collected for the control and treated samples to an approximate depth of 15X coverage to assess the genomic impact of elevated ammonia exposure. Variant discovery identified a total of 79,097 indels and 310,597 SNPs (Supporting Tables S1 and S8 respectively). Of the 310,597 SNPs, we found a total of 4,785 SNPs that were contained within protein coding or regulatory sequences including: untranslated regions (UTRs), exons, and introns (Supporting Table S9). A PANTHER analysis of SNPs was conducted and identified 1,806 coding genes impacted in the ammonia-stressed cultures. The molecular functions of the affected genes and their biological processes are summarized in Fig 3. Binding and catalytic activity are the top 2 molecular function categories, accounting for 35.1% and 34.6% respectively. The other 5 major impacted molecular function categories include transporter activity (8.15%), molecular function regulator (6.62%), transcription regulator activity (6.04%), molecular transducer activity (5.67%), and structural molecule activity (3.56%). The major biological processes impacted by SNP genes involve cellular process (32.7%), metabolic process (18.8%), biological regulation (15.3%), localization (11.3%), and multicellular organismal process (7.88%).

**Fig 3:**
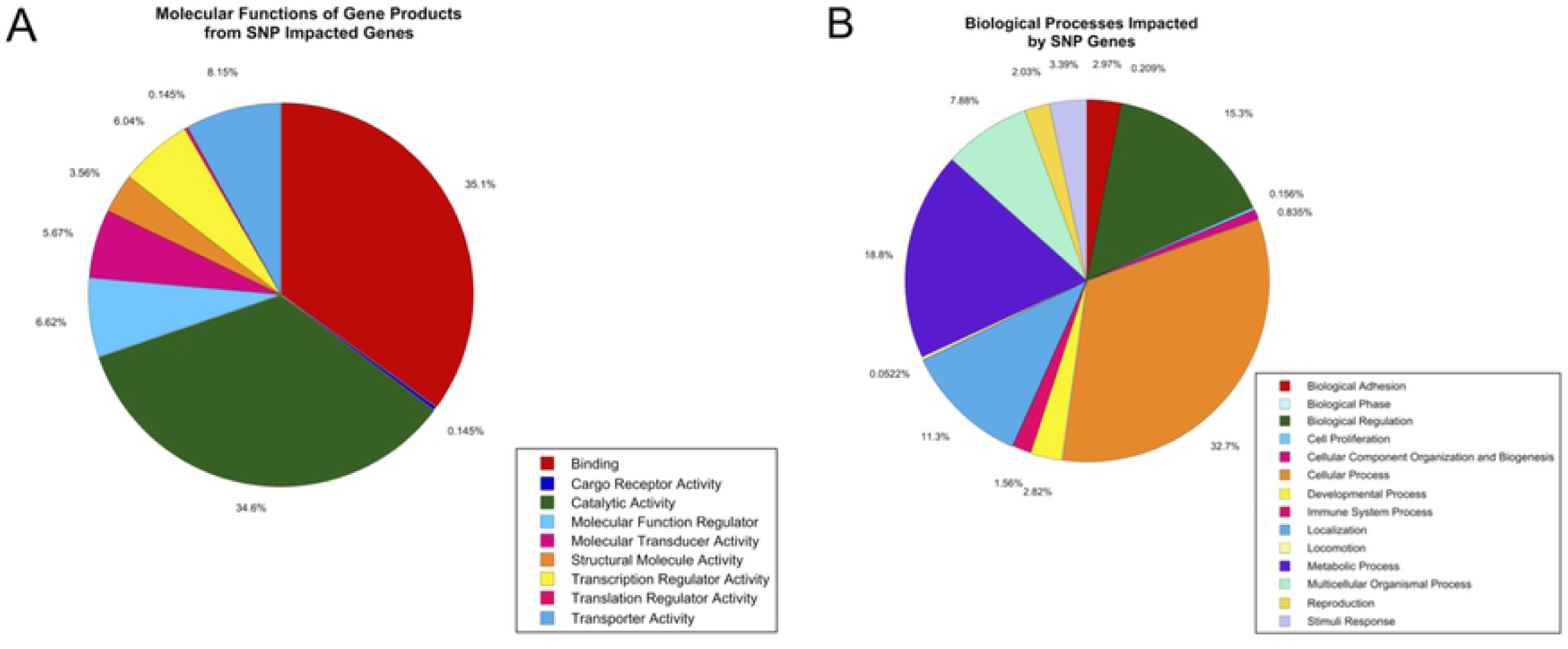
Classification of the 1,806 SNP-impacted genes by A) molecular function of gene product and B) biological processes disturbed.

### Functional enrichment profiling of genes with high-impact SNPs

As there are significantly more literature reports on genome instability in human than in Chinese hamster, a text and data mining approach was used to identify potential Chinese hamster genes involved in genome instability under ammonia stress by use of orthology to human. Through text/data mining, 3,054 human genes were identified as candidate genes related to genome instability by the PubMed search with “genomic instability [MeSH Terms]”, combined with PubTator gene annotations (Supporting Table S15). Among them, 2,521 genes had 1-to-1 protein-coding orthologs in Chinese hamster genome (Supporting Table S15). DAVID gene enrichment analysis identified highly enriched GO-Biological Process (BP), GO-Molecular Function (MF) and KEGG-pathway among these 2,521 human genes. Hundreds of genes were enriched in DNA repair, DNA damage response, DNA replication, cell division, transcription regulation and apoptotic process regulation (Supporting Table S16).

For the 1276 Chinese Hamster genes with detected SNPs of high (e.g., stop gained, splice donor variant and intron variant) and moderate (e.g., missense variant and splice region variant) effect in our ammonia stressed samples, the corresponding human orthologs were subject to DAVID analysis to identify enriched functions (Supporting Tables S11-S14). Top enriched KEGG pathways include Glycosaminoglycan biosynthesis, ECM-receptor interaction, ribosome biogenesis, focal adhesion, Amoebiasis, NOD-like receptor signaling pathway, and regulation of actin cytoskeleton (Supporting Table S13).

For the 122 genes with high SNP effect, DAVID generated gene clusters and functional annotation clusters mostly with relatively low enrichment scores, which suggests quite distinct functions among those genes. Table 1 lists the identified genomic instability (GI) genes by mapping with text-mined human GI genes, as well as referencing the review [41]. It is notable that DAVID identified four genes in pyrimidine metabolism (KEGG pathway hsa00240), three genes in DNA replication (KEGG pathway hsa03030), four genes in carbohydrate metabolic process (GO:0005975), three genes in telomere maintenance via recombination (GO:0000722), five genes in mitotic nuclear division (GO:0007067), and four genes in negative regulation of neuron apoptotic process (GO:0043524) (see details in Supporting Table S10). To check against the results from DAVID, especially mindful of its knowledgebase last updated in 2016, GO enrichment analysis was also conducted on http://geneontology.org/ with tool powered by PANTHER Classification System [42], which is maintained up to date with GO annotations (Supporting Tables S11 and S13). It is affirmative that over twenty genes are identified as involved in cell cycle (GO:0007049) by PANTHER enrichment analysis (Table 1 and Supporting Table S10).

**Table 1:** Genome instability genes with high SNP effect detected in ammonia-stressed samples.

| Chinese Hamster Entrez gene ID | Human Entrez gene ID | Human Gene Name | Function | PMID<sup>c</sup> |
| --- | --- | --- | --- | --- |
| 100761842 | 1763 | DNA replication helicase/nuclease 2(DNA2) | Replication, Cell cycle<sup>d</sup> | 18702299; 20131965; 22960744; 22987153 |
| 100772412 | 6117 | replication protein A1(RPA1) | Replication, Cell cycle<sup>d</sup> | 16523685; 18283122; 18449195; 20103649 |
| 100755630 | 1107 | chromodomain helicase DNA binding protein 3(CHD3) | Chromatin, Cell cycle<sup>d</sup> | 21447119 |
| 100771321 | 23514 | scaffolding protein involved in DNA repair(SPIDR) | Double-strand break repair | 23603433 |
| 100766371 | 64210 | MMS19 homolog, cytosolic iron-sulfur assembly component(MMS19) | DNA repair | 22678362 |
| 100751914 | 4299 | AF4/FMR2 family member 1(AFF1) | Transcription factor activity | 18930318; 8824884; 9299237 |
| 100761435 | 10765 | lysine demethylase 5B(KDM5B) | Transcription factor activity | 24778210; 25714495 |
| 100767456 | 2113 | ETS proto-oncogene 1, transcription factor(ETS1) | Cell cycle, transcription factor activity | 2527802; 28301528; 3422214 |
| 100753037 | 80184 | centrosomal protein 290(CEP290) | Cell cycle | 22797915; 24816561 |
| 100757441 | 54821 | ERCC excision repair 6 like, spindle assembly checkpoint helicase(ERCC6L) | Cell cycle | 26256213; 26643143 |
| 100758610 | 23552 | cyclin dependent kinase 20(CDK20) | Cell cycle | 16413100 |
| 100761529 | 57521 | regulatory associated protein of MTOR complex 1(RPTOR) | Cell cycle | 20013896; 24906312 |
| 100761741 | 10763 | nestin(NES) | Cell cycle | 19216795; 20945392 |
| 100763340 | 9821 | RB1 inducible coiled-coil 1(RB1CC1) | Cell cycle | 19888451 |
| 100763926 | 5727 | patched 1(PTCH1) | Cell cycle | 17204027; 18955550; 19164512; 21699520 |
| 100770459 | 53632 | protein kinase AMP-activated non-catalytic subunit gamma 3(PRKAG3) | Cell cycle | 17693186 |
| 100772150 | 4040 | LDL receptor related protein 6(LRP6) | Cell cycle | 26103566 |
| 100774506 | 899 | cyclin F(CCNF) | Cell cycle | 22632967; 22661117;<br>23182110 |
| 100750774 | 2187 | Fanconi anemia complementation group B(FANCB) | Others | 1581882; 16675878;<br>16803569; 3856472 |
| 100751960 | 4953 | ornithine decarboxylase 1(ODC1) | Others | 18474008 |
| 100753024 | 257194 | neuronal growth regulator 1(NEGR1) | Others | 28235956 |
| 100753715 | 11169 | WD repeat and HMG-box DNA binding protein 1(WDHD1) | Others, Cell cycle<sup>d</sup> | 17222391; 23584157;<br>27099318; 28815832 |
| 100757331 | 1362 | carboxypeptidase D(CPD) | Others | 23460817 |
| 100760540 | 4316 | matrix metalloproteinase 7(MMP7) | Others | 15668893; 20372795;<br>23682078 |
| 100760694 | 10752 | cell adhesion molecule L1 like(CHL1) | Others | 17222391; 19099189;<br>21968058; 24475022 |
| 100763919 | 221981 | thrombospondin type 1 domain containing 7A(THSD7A) | Others | 20804921 |
| 100764526 | 50506 | dual oxidase 2(DUOX2) | Others | 30318012 |
| 100765379 | 5922 | RAS p21 protein activator 2(RASA2) | Others | 26925973 |
| 100765803 | 3927 | LIM and SH3 protein 1(LASP1) | Others | 19688977 |
| 100766952 | 4619 | myosin heavy chain 1(MYH1) | Others | 28696027 |
| 100773500 | 6000 | regulator of G-protein signaling 7(RGS7) | Others | 14623461 |
| 100774312 | 214 | activated leukocyte cell adhesion molecule(ALCAM) | Others | 21935604; 23060121;<br>23065858; 27494849 |
| 100772954 | 5558<sup>a</sup> | primase (DNA) subunit 2(PRIM2) | Replication, Cell cycle<sup>d</sup> |  |
| 100774671 | 5426<sup>b</sup> | DNA polymerase epsilon, catalytic subunit(POLE) | Replication | 25948378 |
| 100753096 | 5981<sup>b</sup> | replication factor C subunit 1(RFC1) | Replication | 15725622; 16523685;<br>20227372; 20573375 |
| 100757579 | 23309<sup>a, b</sup> | SIN3 transcription regulator family member B(SIN3B) | Chromatin |  |
| 100762862 | 641<sup>b</sup> | Bloom syndrome RecQ like helicase(BLM) | Double-strand break repair | 11138010; 11640873;<br>14695165; 15803320 |
| 100770724 | 672<sup>b</sup> | BRCA1, DNA repair associated(BRCA1) | Double-strand break repair | 11832208; 14559796;<br>15162061; 15254237 |
| 100689331 | 5893<sup>b</sup> | RAD52 homolog, DNA repair protein(RAD52) | Double-strand break repair | 15589151; 17222391;<br>17405295; 19112184 |
| 100773161 | 7919<sup>a, b</sup> | DExD-box helicase 39B(DDX39B) | mRNA biogenesis |  |
| 100761220 | 2175<sup>b</sup> | Fanconi anemia complementation group<br>A(FANCA) | Others | 10468606; 10921325;<br>11640873; 16676573 |
| 100758700 | 23137<sup>b</sup> | structural maintenance of chromosomes 5(SMC5) | Others | 16892052; 19217832;<br>21415594; 21550342 |

a. These genes were mapped from 2013 review paper [41], not found in the text mined genomic instability gene list.
b. These genes were mapped from 2013 review paper [41], but identified with moderate SNP effect from our ammonia stressed samples.
c. PMIDs for the text mined Genomic Instability human genes, a maximum of four PMIDs shown.
d. Additional genes identified with cell cycle (GO:0007049) enrichment via Panther enrichment analysis [42].

### Microsatellite and candidate MSI loci

A whole-genome scan for microsatellites discovered a total of 409,628 loci, with motifs that varied from di- to tetranucleotide repeats (Supporting Table S2). As expected, the microsatellites comprised of dinucleotide repeats were the most prevalent with a total of 287,124. Trinucleotide and tetranucleotide motifs were less abundant with 46,602 and 75,902 occurrences, respectively. From the genomic variant data, 79,097 indels and 310,597 SNPs were observed. Complete lists of indels and SNPs can be found in Supporting Tables S1 and S8 respectively. An ideal locus to be used as a biomarker, demonstrated in Fig 4, would have no mutations in the control cultures, some indel events in 10 mM ammonia-stressed cultures, and more pronounced indel events in 30 mM ammonia-stressed cultures.

**Fig 4:**
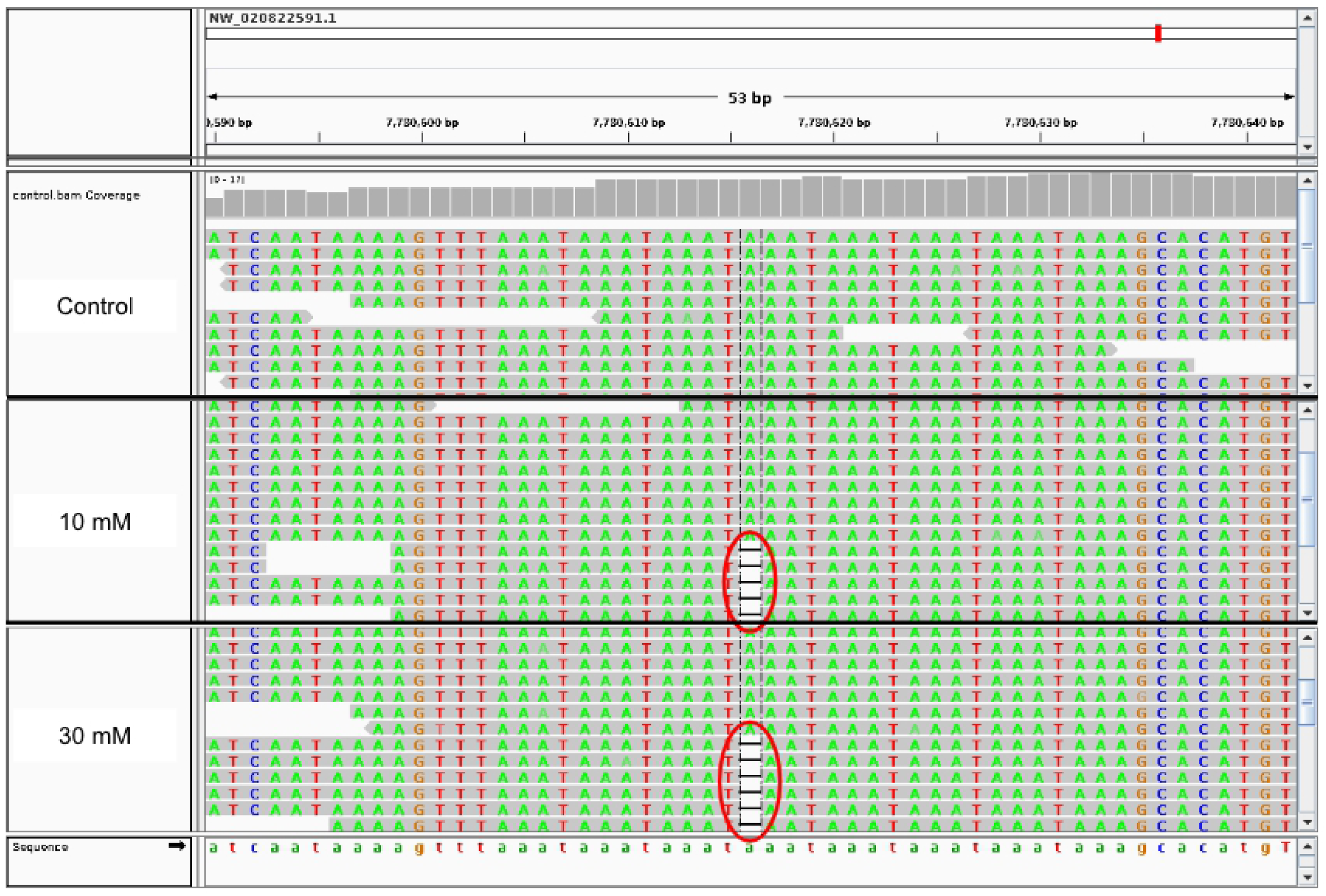
IGV image of a microsatellite located at NW_020822591.1:7780616. Note that the control sample has no mutations, the mildly-stressed sample has five, and the highly-stressed sample has six.

Using a mutation score and stringent filters, we developed a panel set of 124 MSI loci that could be ordered by priority for follow up research. Assigning mutation scores to individual loci was the most effective way to quantify respective mutation rates. It is important to note that because the mutation score is calculated using the allelic depth; loci with more reads are statistically more significant than those with fewer reads. With this in mind, the 124 candidate loci may not be all inclusive of the optimal loci due to the variation in read depth across the genome. The number of loci remaining after each filter step is summarized in Table 2, the full list of loci in each step can be found in Supporting Tables S4-S7, and the location of all candidate loci are summarized in Table 3.

**Table 2:** Numerical representation of filter progression.

| Filter Criteria | Remaining Loci |
| --- | --- |
| Genome-wide INDELs | 79,097 |
| 30 mM Relative Mutation Score > 10 mM Relative Mutation Score | 35,437 |
| Loci with positive Relative Mutation Score | 16,678 |
| Loci confirmed to be microsatellites | 187 |
| Loci with genotypes that vary from the control | 124 |

**Table 3:** Location and composition of all candidate microsatellites. More detailed information on candidate loci can be found in Supporting Table S7.

| Scaffold | Position | Motif | Mutation Scores (%) |  |  |
| --- | --- | --- | --- | --- | --- |
|  |  |  | 10 mM | 30 mM | Control |
| NW_020822370.1 | 34498325 | (GA)31 | 85.71 | 94.44 | 83.33 |
| NW_020822370.1 | 29116483 | (AG)22 | 14.29 | 62.5 | 0 |
| NW_020822370.1 | 8110802 | (GT)33 | 14.29 | 20 | 7.14 |
| NW_020822373.1 | 4520712 | (TC)27 | 10.34 | 12.4 | 4.59 |
| NW_020822375.1 | 17984519 | (GT)23 | 90 | 100 | 77.78 |
| NW_020822375.1 | 24391224 | (CA)8 | 41.67 | 70 | 0 |
| NW_020822376.1 | 2957094 | (CA)7 | 13.04 | 21.05 | 8.33 |
| NW_020822382.1 | 3087333 | (CCTC)5 | 14.29 | 21.43 | 0 |
| NW_020822403.1 | 19387296 | (AC)25 | 28.57 | 50 | 4.76 |
| NW_020822403.1 | 15933029 | (AC)26 | 25 | 46.15 | 0 |
| NW_020822406.1 | 3701096 | (GT)23 | 14.29 | 42.86 | 9.09 |
| NW_020822407.1 | 7711795 | (AAAC)6 | 16.67 | 26.67 | 0 |
| NW_020822409.1 | 5651224 | (AAT)12 | 14.29 | 44.44 | 0 |
| NW_020822410.1 | 13888699 | (GT)28 | 83.33 | 90.32 | 63.64 |
| NW_020822410.1 | 10858329 | (AC)19 | 94.12 | 96.15 | 75 |
| NW_020822412.1 | 3775426 | (TG)26 | 21.43 | 28.57 | 0 |
| NW_020822415.1 | 36220614 | (TC)31 | 91.67 | 92.31 | 57.14 |
| NW_020822415.1 | 16052939 | (ACA)5 | 19.23 | 23.33 | 5.26 |
| NW_020822421.1 | 5779917 | (TC)29 | 91.3 | 100 | 80 |
| NW_020822426.1 | 5559446 | (GA)31 | 93.75 | 100 | 90.91 |
| NW_020822426.1 | 3827444 | (TCT)30 | 71.43 | 72.73 | 55.56 |
| NW_020822426.1 | 1539468 | (GT)22 | 27.27 | 28.57 | 0 |
| NW_020822428.1 | 1367872 | (AC)26 | 93.75 | 100 | 85.71 |
| NW_020822428.1 | 8085398 | (TCCT)11 | 16.67 | 27.27 | 0 |
| NW_020822434.1 | 512274 | (GT)28 | 18.18 | 44.44 | 0 |
| NW_020822436.1 | 3837117 | (AC)16 | 90.32 | 93.33 | 38 |
| NW_020822439.1 | 68421902 | (AC)29 | 91.67 | 100 | 83.33 |
| NW_020822439.1 | 47065858 | (TC)35 | 84.62 | 94.74 | 80 |
| NW_020822439.1 | 62174741 | (GT)23 | 22.22 | 35.48 | 11.11 |
| NW_020822439.1 | 14184660 | (AAAC)7 | 12 | 50 | 0 |
| NW_020822439.1 | 66094811 | (AC)24 | 25 | 28.57 | 0 |
| NW_020822439.1 | 16584719 | (GT)8 | 20 | 36.36 | 7.14 |
| NW_020822439.1 | 2390672 | (TG)62 | 12.5 | 33.33 | 0 |
| NW_020822439.1 | 243530 | (CA)25 | 25 | 50 | 0 |
| NW_020822439.1 | 17884454 | (AGAT)11 | 13.33 | 20 | 7.14 |
| NW_020822440.1 | 3560698 | (GA)28 | 88.89 | 90 | 83.33 |
| NW_020822443.1 | 2115154 | (AC)27 | 27.27 | 43.75 | 9.09 |
| NW_020822446.1 | 382267 | (CA)32 | 66.67 | 75 | 33 |
| NW_020822450.1 | 9390312 | (AC)22 | 16.67 | 44.44 | 11.11 |
| NW_020822452.1 | 9210374 | (CT)31 | 90 | 93.33 | 81.82 |
| NW_020822458.1 | 15628677 | (AG)27 | 91.67 | 100 | 73.33 |
| NW_020822461.1 | 21799265 | (TCTT)16 | 16.67 | 25 | 0 |
| NW_020822464.1 | 1598541 | (AC)26 | 84.62 | 100 | 83.33 |
| NW_020822464.1 | 6242934 | (TG)26 | 27.78 | 36 | 0 |
| NW_020822465.1 | 1645147 | (TG)9 | 39.29 | 54.55 | 5.88 |
| NW_020822468.1 | 15157395 | (GT)35 | 87.5 | 100 | 66.67 |
| NW_020822469.1 | 3552281 | (GATG)8 | 31.25 | 54.55 | 0 |
| NW_020822484.1 | 1703831 | (TG)7 | 95.65 | 100 | 77.27 |
| NW_020822486.1 | 16687223 | (GT)23 | 21.43 | 22.22 | 0 |
| NW_020822487.1 | 21884245 | (CT)34 | 89.66 | 96.43 | 80 |
| NW_020822487.1 | 17786455 | (TC)30 | 91.67 | 100 | 87.5 |
| NW_020822487.1 | 25648949 | (AC)15 | 25 | 37.5 | 10 |
| NW_020822487.1 | 8954697 | (CA)29 | 10 | 18.18 | 0 |
| NW_020822488.1 | 1550207 | (TG)26 | 85.71 | 100 | 75 |
| NW_020822499.1 | 2670448 | (TG)31 | 95.65 | 100 | 84 |
| NW_020822499.1 | 21810339 | (CT)20 | 35.71 | 56.25 | 0 |
| NW_020822499.1 | 16894388 | (TTA)5 | 11.9 | 13.73 | 6.67 |
| NW_020822499.1 | 10821740 | (TTGT)8 | 14.29 | 38.46 | 7.14 |
| NW_020822501.1 | 11357080 | (GT)29 | 91.67 | 100 | 85.71 |
| NW_020822501.1 | 14044688 | (TTG)7 | 14.29 | 24.32 | 3.33 |
| NW_020822501.1 | 14071142 | (TTA)12 | 20 | 28.57 | 5.56 |
| NW_020822501.1 | 17408028 | (TG)30 | 55.56 | 57.14 | 7.14 |
| NW_020822503.1 | 10248411 | (CTTT)14 | 22.22 | 25 | 0 |
| NW_020822503.1 | 4513778 | (GT)14 | 10.53 | 20 | 0 |
| NW_020822503.1 | 17353421 | (GT)6 | 22.22 | 25 | 9.09 |
| NW_020822504.1 | 9433369 | (TC)30 | 95 | 100 | 88.89 |
| NW_020822504.1 | 13319850 | (TGTA)5 | 12.5 | 15.56 | 5.88 |
| NW_020822505.1 | 16324767 | (CA)24 | 93.75 | 100 | 77.78 |
| NW_020822505.1 | 16510213 | (AC)30 | 85.71 | 87.5 | 77.78 |
| NW_020822505.1 | 10642584 | (TG)27 | 27.27 | 37.5 | 0 |
| NW_020822508.1 | 1001369 | (AC)22 | 90.91 | 100 | 71.43 |
| NW_020822508.1 | 2728629 | (TATC)10 | 16.67 | 27.78 | 0 |
| NW_020822508.1 | 15865728 | (GAAG)13 | 7.69 | 31.25 | 0 |
| NW_020822508.1 | 15865731 | (GAAG)13 | 7.14 | 26.32 | 0 |
| NW_020822511.1 | 9568885 | (TG)29 | 92.31 | 100 | 83.33 |
| NW_020822512.1 | 6724926 | (TG)30 | 87.5 | 93.33 | 85.71 |
| NW_020822519.1 | 6274345 | (AC)25 | 91.67 | 100 | 84.62 |
| NW_020822519.1 | 11516391 | (AG)36 | 15.79 | 37.5 | 0 |
| NW_020822520.1 | 2511201 | (AC)24 | 20 | 25 | 0 |
| NW_020822526.1 | 18794354 | (AC)26 | 91.67 | 100 | 83.33 |
| NW_020822526.1 | 25646840 | (CA)7 | 90 | 100 | 77.78 |
| NW_020822526.1 | 24206438 | (GT)30 | 87.5 | 90 | 70 |
| NW_020822526.1 | 16873752 | (AG)32 | 83.33 | 100 | 69.23 |
| NW_020822526.1 | 17494091 | (AC)16 | 17.65 | 57.14 | 0 |
| NW_020822526.1 | 4205760 | (ATGT)11 | 16.67 | 37.5 | 0 |
| NW_020822529.1 | 28253360 | (TG)23 | 91.67 | 100 | 86.67 |
| NW_020822529.1 | 16865510 | (TC)27 | 42.86 | 54.55 | 0 |
| NW_020822530.1 | 10618047 | (TG)22 | 18.18 | 28.57 | 0 |
| NW_020822531.1 | 6287546 | (AC)27 | 90.91 | 93.75 | 81.82 |
| NW_020822531.1 | 1037419 | (GT)44 | 45.45 | 50 | 12.5 |
| NW_020822544.1 | 4160116 | (AAC)5 | 22.73 | 60 | 0 |
| NW_020822567.1 | 4418183 | (TG)29 | 91.67 | 100 | 80 |
| NW_020822567.1 | 6942408 | (AC)24 | 25 | 35.29 | 0 |
| NW_020822567.1 | 3062112 | (CA)7 | 17.39 | 29.41 | 3.85 |
| NW_020822567.1 | 1362729 | (GA)24 | 15.38 | 25 | 0 |
| NW_020822570.1 | 21823495 | (TG)28 | 14.29 | 44.44 | 0 |
| NW_020822591.1 | 7123790 | (TATT)6 | 94.44 | 100 | 80 |
| NW_020822591.1 | 7780615 | (TAAA)8 | 33.33 | 37.5 | 0 |
| NW_020822591.1 | 14941 | (GA)34 | 9.09 | 25 | 0 |
| NW_020822591.1 | 8820175 | (TTAT)11 | 9.52 | 27.27 | 0 |
| NW_020822592.1 | 4061125 | (AC)31 | 42.86 | 44.44 | 0 |
| NW_020822595.1 | 5415636 | (CAAA)6 | 37.93 | 38.46 | 7.69 |
| NW_020822597.1 | 5771854 | (TG)31 | 88.24 | 94.44 | 75 |
| NW_020822601.1 | 72176322 | (CA)8 | 91.18 | 100 | 85.71 |
| NW_020822601.1 | 61385941 | (AC)19 | 21.43 | 22.73 | 0 |
| NW_020822602.1 | 6751043 | (TC)30 | 91.67 | 96.55 | 83.33 |
| NW_020822602.1 | 9765422 | (AATA)7 | 15.38 | 15.79 | 0 |
| NW_020822603.1 | 9112470 | (AC)25 | 92.31 | 100 | 66.67 |
| NW_020822604.1 | 8845685 | (GT)6 | 10 | 37.5 | 0 |
| NW_020822610.1 | 821820 | (AC)27 | 31.25 | 41.18 | 10 |
| NW_020822614.1 | 8982333 | (ATAG)16 | 13.64 | 13.79 | 0 |
| NW_020822629.1 | 5127703 | (AG)28 | 12.5 | 20 | 0 |
| NW_020822629.1 | 6170860 | (CA)22 | 12.5 | 33.33 | 0 |
| NW_020822634.1 | 2791562 | (TC)33 | 95.24 | 100 | 62.5 |
| NW_020822638.1 | 3030009 | (AC)21 | 92.31 | 100 | 58.33 |
| NW_020822698.1 | 6755673 | (GT)27 | 90 | 100 | 80 |
| NW_020822698.1 | 121373 | (TG)30 | 28.57 | 37.5 | 0 |
| NW_020822785.1 | 9557 | (CA)14 | 14.29 | 75 | 0 |
| NW_020822967.1 | 23702 | (CA)17 | 15.38 | 33.33 | 9.09 |
| NW_020823044.1 | 10660 | (TTG)6 | 15.29 | 15.45 | 2.53 |
| NW_020823531.1 | 97528 | (AG)22 | 11.76 | 15.22 | 5.41 |
| NW_020823768.1 | 45602 | (TC)26 | 93.75 | 95.83 | 78.95 |
| NW_020824031.1 | 35819 | (AC)23 | 20 | 25 | 0 |
| NW_020824065.1 | 30045 | (AG)6 | 12 | 14.29 | 4.65 |

## Discussion

Ammonia is a common metabolic waste product in cell culture, and the accumulation of this waste leads to decreased productivity of the culture over time. In this study, mild (10 mM) and high (30 mM) ammonia stresses were used to investigate the effects of this byproduct on genome stability. The VCD, cell viability, and metabolic profiles indicated that the high ammonia stress significantly impacted the culture health, as most of these profiles were significantly different from the control and mild ammonia stress culture within 96 hours (4 days) of the ammonia addition (Fig 2). Additionally, the product titer for the high ammonia stressed culture was severely lowered (Fig 2C). In contrast, the culture profiles for the mild ammonia stressed cultures were not significantly different from the control, except for the alanine profile beginning at 84 hours (Day 3.5). However, the effects of the mild stress were eventually realized around the 132 hour point (Day 5.5) where we observed reduced productivity. These culture profiles indicate that ammonia impacts CHO cells in a dose dependent manner, and even mild ammonia stress has an influence on culture productivity. Until now, efforts have mainly focused on transcriptome and proteome effects of ammonia stress [11, 43, 44].

The relatively abrupt effects of ammonia on culture health and productivity can be linked to genetic underpinnings. Whole genome sequencing and variant discovery of SNPs allowed for the identification of genotypic changes caused by the ammonia stress. More than 1800 gene variants were identified to be unique in the ammonia stressed samples, including genes involved in MMR. An impaired or inefficient MMR system can lead to the accumulation of mutations in functional genes over cell divisions that are critical to the cell’s survival and can lead to loss of genetic stability [45] or disease states, such as cancer [46]. The need for a highly conserved MMR system can be observed by the presence of multiple orthologs of *MutS* and *MutL* in eukaryotic genomes [47]. *MutS* binds to base mismatches or small indels [48, 49] while *MutL* is responsible for communicating the identification of mismatch events to downstream elements of MMR such as exonucleases [50].

Under normal circumstances, DNA replication errors are a normal part of the replication process, but are usually corrected immediately through proofreading or mismatch repair [51]; however, exposure to environmental stress has been shown to underlie error correction fidelity of the MMR machinery, leading to an accelerated rate of variant accumulation and loss of genetic stability [52]. In this study, ammonia stress led to the accumulation of genomic variants, which were observed within 72 hours of the stress (culture time 84 hours). Variant effect prediction found high-impact mutations [26], such as loss of function, frame shift, stop loss, and start gain that leads to an altered gene function in genes involved in genome stability, as a result of ammonia stress (Table 1). For example, critical genes, such as Breast Cancer susceptibility gene 1 (*BRCA1*), *RAD52*, DNA Replication Helicase/Nuclease 2 (*DNA2*), and DNA Polymerase Epsilon, Catalytic Subunit (*POLE*) were impaired. *BRCA1* is a tumor suppressor gene that serves a variety of cellular processes including DNA repair, maintaining chromosome stability, and transcription regulation in response to DNA damage. Loss of function mutations in *BRCA1* have been linked to familial breast and ovarian cancer due to the critical role it plays in genome stability [53]. Many studies of *RAD52* in yeast have identified its role in homology directed repair as a recombination mediator that allows for the repair of double-strand breaks [54-56]. Without homology directed repair, double-strand breaks are considerably more likely to be repaired inaccurately. Furthermore, a recent study suggests that *RAD52* also works to stabilize stalled replication forks and prevent their collapse [57].

Other biological processes and molecular functions that were affected included DNA integrity maintenance, carbohydrate metabolic processes, and pyrimidine metabolism. Genes such as alpha 1,3-galactosyltransferase 2 (*A3GALT2*), carbohydrate sulfotransferase 3 (*CHST3*), glutamine-fructose-6-phosphate transaminase 1 (*GFPT1*), glycerol kinase 2 (*GK2*) had variants, where the reading frame was altered which could affect gene function or result in complete loss of function. Additionally, some moderate impact variants affected genes with functional roles in other biological processes and pathways that might contribute to recombinant protein expression. One example of an observed moderate impact variant is impaired ribosome biogenesis, which could be correlated with the drastic difference in titers of the cultures with different levels of ammonia stress.

Variants in genes that regulate the MMR pathway may be an origin to the cascade of events that leads to genome instability. When the MMR pathway in a cell is compromised, mistakes can occur and propagate indiscriminately across the genome as cell division occurs [47-49] Unfaithful replication of genomic repeats, such as microsatellite repeats, have been used as effective biomarkers in predicting certain diseases, such as cancer [16]. For example, the Bethesda Panel is comprised of only 5 microsatellites in the human genome. The diagnostic screen amplifies and sequences this stable set of microsatellite loci, and if more than one locus exhibits indel mutations, the patient is deemed MSI high, which is indicative of their cancer being attributed to genome instability [21]. Moreover, the fact that only five MSI loci are used for the diagnostic demonstrates how few microsatellite loci in the human genome exhibit the ideal behavior of a diagnostic biomarker. In the analysis of the CHO genome under ammonia stress, more than 400,000 predicted microsatellites and nearly 80,000 indels were discovered. When intersecting these two datasets, only 1,022 microsatellites were observed to have indels (Supporting Table S3). To identify priority and rank the list of candidate MSI’s for possible use as a diagnostic tool for CHO cell instability, a ranking strategy was developed that accounted for read depth and mutation frequency, as determined by division of the total number of variant reads by total number of reads. This ranking strategy resulted in a list of 124 candidate loci (Table 3) with the potential to be reliable biomarkers of genome instability for CHO cell cultures.

For this study, the genome was sequenced to a resolution of 15 reads per base pair. In some regions, the allelic sequencing depth was much higher than others, which could impact the mutation score method employed in this procedure. Nonetheless, the employed filters produced a sufficient number of candidate loci for further experimentation. Indels identified within candidate loci had dose dependent responses to the ammonia concentrations that were not observed in the control cultures. Though our filtering criteria identified 124 candidate loci that have the potential to be used on a diagnostic panel, they should be amplified and sequenced individually in order to form a concise and reliable panel similar to that of the Bethesda Panel.

## Conclusion

The accumulation of metabolic wastes, such as ammonia, has profound effects on CHO culture viability and product quality. Past work has tied ammonia stress to changes in the transcriptome as well as product quality. In this study, the genotoxicity of ammonia was investigated for mild and high ammonia stresses. Both stresses caused observable culture changes, where the high ammonia stress had observable effects on VCD and cell viability soon after the introduction. In contrast, the mild ammonia-stressed culture was not significantly impacted, with the exception of certain metabolites, such as alanine. Yet, the titer production was lower than expected for the mild ammonia-stressed cultures. Interestingly, genome analysis identified many SNPs and indels as well as some potential MSI loci in the mild and high ammonia-stressed cultures, where these mutations were observed to have a dose dependent response. These mutations are propagated through the culture as cells divide and have detrimental impacts on metabolism, reproduction, signaling pathways, as well as many other cellular mechanisms. Furthermore, we have demonstrated that a sufficient number of candidate MSI loci exist within the CHO genome that could be used as DNA biomarkers to monitor culture genomic health. The development of a simple MSI diagnostic could assist in assessing the degree of genome instability in cultures prior to metabolite shifts, which would lead to precision-driven culture system that would ultimately increase productivity. Future applications will be to determine which of the presented candidate MSI loci would be the most effective in a diagnostic panel and to determine whether or not they are stable in other common culture stresses.

## Supporting Table Captions

**Supporting Table S1:** All reported indels across the CHO genome as a result of ammonia stress.

**Supporting Table S2:** The genome-wide MISA report that predicted all microsatellite regions in the CHO genome.

**Supporting Table S3:** The intersection of tables S1 and S1 that identifies all microsatellites that contain indels.

**Supporting Table S4:** Loci remaining after the first filtering step.

**Supporting Table S5:** Loci remaining after the second filtering step.

**Supporting Table S6:** Results of a MISA report used to find microsatellites within the reported indels from table S1

**Supporting Table S7:** All candidate loci identified as potential MSI biomarkers indicative of genome instability.

**Supporting Table S8:** All reported SNPs across the CHO genome as a result of ammonia stress.

**Supporting Table S9:** SNPs from table S8 that were within coding or regulatory regions.

**Supporting Table S10:** High-impact SNPs that contain human orthologs to genes involved in maintaining genome stability.

**Supporting Table S11:** Results of DAVID enrichment of high-impact SNPs.

**Supporting Table S12:** High-impact SNPs in CHO without a human ortholog.

**Supporting Table S13:** DAVID enrichment of high and moderate-impact SNPs.

**Supporting Table S14:** Panther enrichment of high-impact SNPs.

**Supporting Table S15:** Human genes associated with genome instability.

**Supporting Table S16:** Enrichment of the 2521 human genome instability genes.

